# Precursor microRNA Identification Using Deep Convolutional Neural Networks

**DOI:** 10.1101/414656

**Authors:** Binh Thanh Do, Vladimir Golkov, Göktuğ Erce Gürel, Daniel Cremers

## Abstract

Precursor microRNA (pre-miRNA) identification is the basis for identifying microRNAs (miRNAs), which have important roles in post-transcriptional regulation of gene expression. In this paper, we propose a deep learning method to identify whether a small non-coding RNA sequence is a pre-miRNA or not. We outperform state-of-the-art methods on three benchmark datasets, namely the human, cross-species, and new datasets. The key of our method is to use a matrix representation of predicted secondary structure as input to a 2D convolutional network. The neural network extracts optimized features automatically instead of using a large number of handcrafted features as most existing methods do. Code and results are available at https://github.com/peace195/miRNA-identification-conv2D.

## 1. Introduction

Ribonucleic acids (RNAs) are biomolecules involved in many biological processes. RNA is assembled as a chain of monomers called nucleotides. There are four types of RNA nucleotides that serve as arbitrarily arranged building blocks of the nucleotide chain: *adenine* (A), *cytosine* (C), *guanine* (G), and *uracil* (U). MicroRNAs (miRNAs) are a type of non-coding RNA molecule, usually about 20-23 nucleotides long. Over the last three decades, they have been detected in a large number of organisms such as humans [5] and plants [23], as well as in viruses [37]. They bind to target messenger RNAs (mRNAs) to inhibit the translation of mRNAs to proteins [3]. Their importance in gene regulation plays a part in diseases such as cancer [20, 10, 46], and they are good targets for disease markers and therapeutics [46]. Thus, miRNAs identification is a crucial task in medi-cal treatments. But it is difficult to identify miRNAs directly because they are short. Most studies focus on computational methods for identifying precursor miRNAs (pre-miRNAs) instead. Primary transcripts called pri-miRNAs are processed to form pre-miRNAs and then to mature miRNAs. Identifying pre-miRNAs is easier in comparison to miRNAs because pre-miRNAs are a lot longer (approximately 80 nucleotides) and they have a hairpin loop structure with more structural features. Pre-miRNA identification is a classification task, yielding the output “positive” or “negative”. It is potentially a hard task due to the enormous amount of possible sequences that can be arranged using 4 nucleotide types A, C, G and U. Moreover, the number of explored pre-miRNAs is much smaller than the number of pseudo hairpins (i.e. RNAs which have similar hairpin loop structure to pre-miRNAs but do not contain mature miRNAs). Hence, we have to cope with a class imbalance problem. On the other hand, RNAs have many structural and biological features and we do not know which ones are really needed for pre-miRNAs identification. Therefore, machine learning is an appropriate approach to weigh features automatically.

Previous work on miRNA and pre-miRNA identification has been based on handcrafted rules (MIReNA [32]) or machine learning. Machine learning methods have been increasingly popular during the last decade and demonstrated to be the most promising, with tools such as HuntMi [16], miRBoost [43], CSHMM [1], microPred [4], miPred [35],triplet-SVM [47], Mirann [38], DP-miRNA [44], deepMiR-Gene [36]. They predict secondary structure (base-pairing interactions within the RNA sequence, see Fig. 1AB for an example) with standard methods such as RNAfold [19], GT-fold [33], and CyloFold [6]. Then they extract numerous handcrafted features, such as counts of Watson–Crick nucleotide pairs (A-U, C-G), loop length [44, 43], sequence length [44], dinucleotide pair frequencies [44, 21, 43, 4, 35], trinucleotide pair frequencies (constituting 64 features) [44, 21], melting temperature [44], minimum free energy [45, 44, 9]. These features are used as inputs to machine learning methods such as support vector machines (SVM) [43, 4, 47], random forests [35], neural networks [38, 44, 36, 21] and hidden Markov models [1]. For example, Ref. [21] uses 98 of the aforementioned features for the input of their neural network. DP-miRNA [44] uses a neural network with 58 extracted characteristic features based on sequence composition and secondary structure predicted by RNAfold software, folding measures including various formulations of physical energy. One of the most feature-rich methods is Ref. [9], which uses around 900 features. The authors showed that the most significant factor in pre-miRNA identification is the secondary structure. Secondary structures are distinctive and many features can be extracted from them. The state-of-the-art method deepMiRGene [36] uses one-hot encoding of so-called dot-bracket notation to represent RNA secondary structure as input to a neural network. The problem with this representation is that the information contained therein is rather “entangled”: There is no single data entry that indicates whether two nucleotides {*i, j}* are paired with each other. Finding out whether they are paired requires parsing much of the entire secondary structure representation. This parsing is achieved by training a neural network with memory and attention mechanisms. However, “outsourcing” such a known meaningful information disentanglement to the learning usually results in suboptimal disentanglement and a more difficult overall learning task [28]. The biggest advantage of deepMiRGene is they do not need any handcrafted features [36].

**Figure 1:**
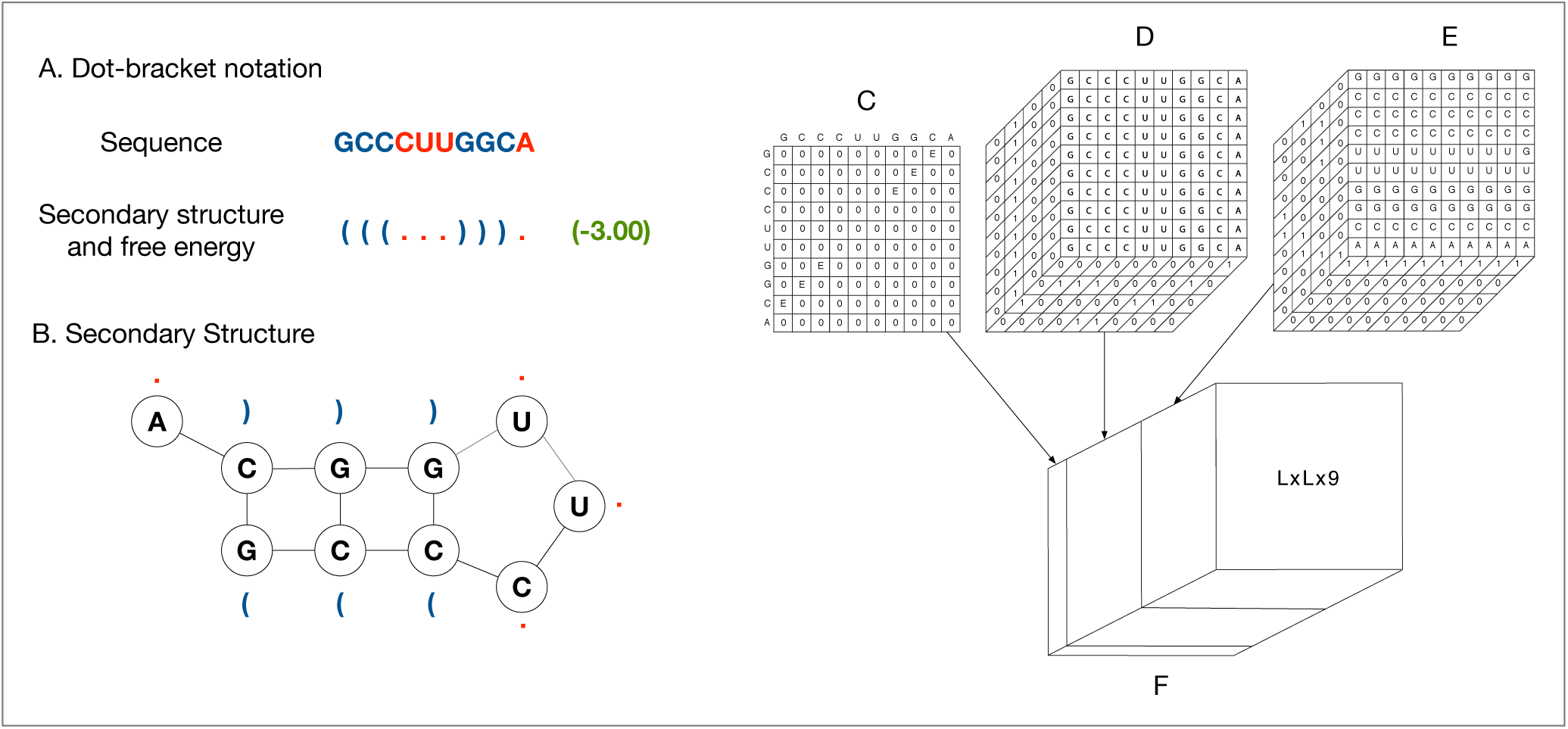
Example for preprocessing step of “GCCCUUGGCA” sequence. (A) Sequence of a pre-miRNA and the dot-bracket energy. In dot-bracket notation, each corresponding “()” structure of the given sequence. (C) is the pairing matrix nimum free energy. (D) and (E) are vertical and horizontal notation of its secondary structure, −3.00 is the minimum of free pair represents an interaction nucleotide pair. (B) is the secondarybased on the secondary structure of sequence where *E* is the mi one-hot encoded sequence. (F) is the combination of (C), (D) and (E) by concatenating.

Deep learning can extract the high-level hidden features of the input very well and get good performance on many tasks such as image classification, face and speech recognition [29], molecular function prediction [34, 14], protein secondary [40] and tertiary structure prediction [2], protein contact prediction [13]. Inspired by the success of deep learning and the importance of secondary structure for pre-miRNA identification, we propose an end-to-end deep learning method using the given input sequence and its secondary structure. For the sequence data, people often use LSTM [18] neural networks because they can handle information that is far away in the sequence effectively. However, in this paper, we are going to use a convolutional neural network (ConvNet) in order to classify RNA sequences as pre-miRNA. We use a downscaling layer to consolidate features and to allow variable-sized input of the ConvNet to lead to a fixed-sized class prediction. The key of our approach is to encode the secondary structure in the pairing matrix format (also known as *dot plot*, see [8, 12, 31, 42]). In the pairing matrix, we specify the minimum free energy of the sequence and the interactions between nucleotides directly. The results show that our algorithm with this new technique performs better than other state-of-the-art methods on the benchmarks, namely the so-called human,cross-species and new datasets [36, 45]. When we compared our results with and without the pairing matrix, we saw that the results are much worse without the pairing matrix. Our contributions are:

- A novel joint 2D multi-channel representation of sequence, secondary structure, and minimum free energy without handcrafted features,
- A convolutional network that is appropriate for that type of input representation and outperforms state-of-the-art methods,
- A comparison between variable-sized and fixed-sized inputs for the ConvNet.

## 2. Our approach

Our approach consists of two steps. Firstly, we represent the given RNA in a 2D multi-channel format based on sequence one-hot encoding and pairing matrix of its predicted secondary structure. Using this input representation, we train a ConvNet to identify pre-miRNA. Our proposed ConvNet is designed to adapt to variable size of inputs by using a downscaling layer between the convolutional and fully connected layers. For comparison, we also build a fixed-sized inputs ConvNet by zero-padding the inputs to have the same size.

### 2.1 Input representation

Secondary structure is a very beneficial feature for pre-miRNA identification [22, 21, 9]. In this paper, we represent predicted secondary structure using a pairing matrix format as given in Fig. 1C, where for every pair of nucleotides 0stands for a non-interaction, and a non-zero value (we use the minimum free energy *E* of the molecule) stands for an interaction.

Each nucleotide A, C, G, U of the input sequence is represented using one-hot encoding, which is a binary vector of dimension 4. Thus, we have the dictionary*{A*→ (1, 0, 0, 0)*, C* → (0, 1, 0, 0)*, G*→ (0, 0, 1, 0)*, U* → (0, 0, 0, 1) *}*, yielding an *L*× 4 array that represents the entire sequence of length *L* in one-hot encoding. Each RNA input sequence of length *L* is represented as follows (see also Fig. 1 for an example):

- Use RNAfold [19] to predict the secondary structure and minimum free energy.
- Represent the predicted secondary structure as a *L* ×*L pairing matrix*, where position (*i, j*) indicates whether the *i*^t h^ nucleotide interacts with the *j*^t h^ nucleotide or not.
- Reshape the *L* × 4 one-hot encoded sequence to *L* ×1 × 4 and replicate it horizontally to shape *L* × *L* × 4.
- Reshape the *L* × 4 one-hot encoded sequence to 1 ×*L* × 4 and replicate it vertically to shape *L* × *L* × 4.
- Concatenate the three aforementioned arrays along the channels dimension (third dimension), yielding *L* × *L* × (1 + 4 + 4).

By using this data representation, we ensure that every 1× 1× 9 “pixel” (*i, j*) of the L× *L*× 9 array contains the entire available information about the nucleotide pair (*i, j*), namely the nucleotide type at position *i* and at position *j* as well as whether the two nucleotides are× paired. Moreover, the neighborhood of that “pixel” contains the entire available information about sequence neighborhoods of positions *i* and *j* and the pairing between these neighborhoods. The presence and arrangement of such neighborhood-patterns is characteristic of certain RNA types. A convolutional network is appropriate for extracting high-level information from such local patterns (as shown for *L*× *L* representations of proteins [13]), and (by using pooling/downscaling layers) consolidating such information across the entire molecule. Fig. 1 is an example of our preprocessing step for the sequence “GCCCUUGGCA”, where Fig. 1F is the output of this step.

### 2.2 Network architecture and training

Our network architecture is shown in Fig. 2. Both architectures contain three convolutional layers followed by two fully connected layers. The reasons for using such an architecture in combination with our input representation are described in Section 2.1. For comparison, as an alternative to variable-sized inputs (Algorithm 1, Fig. 2A), we pad the *L×L×*9 input to 400*×*400*×*9 by zeros to the bottom and to the right before providing it as input to the ConvNet (Algorithm 2, Fig. 2B). For variable-sized inputs (Algorithm 1), because of variable length *L* of the input sequence, we set the batch size to 1 and use a downscaling layer after the convolutional layers to obtain fixed-size features which will be fed into fully connected layers.

**Figure 2:**
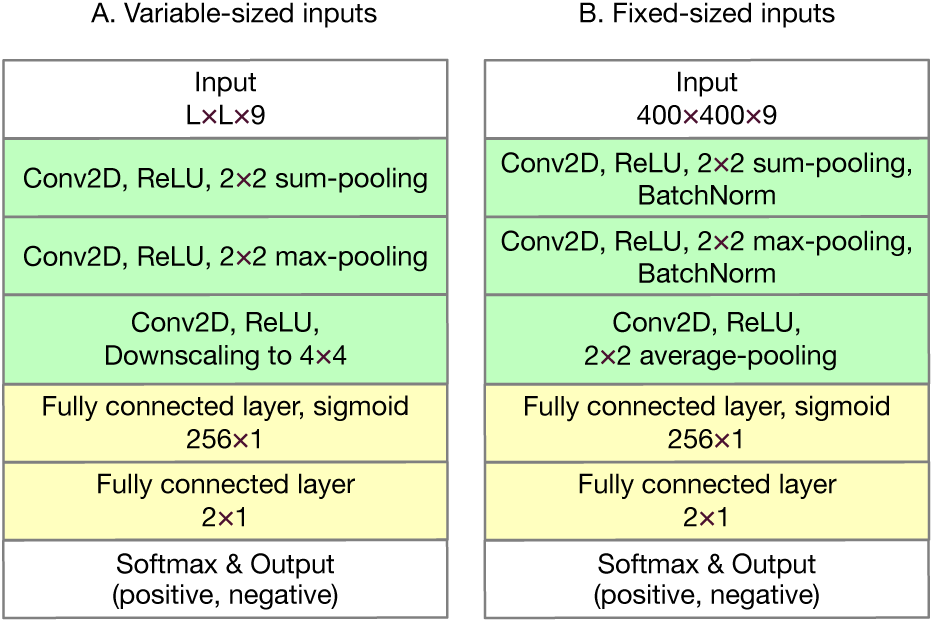
Our proposed ConvNet architecture.

We normalize the outputs to [0, 1] by using a softmax layer at the end. It is calculated as follows, where 2 is the number of labels:

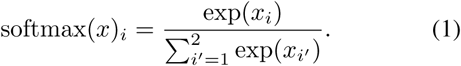

The model is trained by minimizing the cross-entropy loss function:

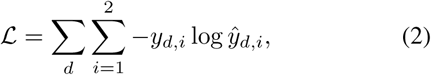

where *d* enumerates training samples, *y*_*d*_ ∈ ℝ ^2^ is (α, 0) if *d* is a positive sample, (0, β) if it is a negative sample(α, β ∈ ℝ are the weights for penalizing class imbalance;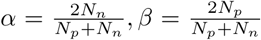 where *N*_*p*_*, N*_*n*_ is the number of positive and negative samples, respectively), and ŷ_*d*_∈ ℝ^2^is the output of the softmax layer.

We trained the network with the adaptive subgradient method (AdaGrad) [11] and a learning rate of 0.001. In alternative experiments, we also tried soft one-hot encoding such as {0.2, 0.8} or {0.1, 0.9} for the input sequenceencoding instead of {0, 1}, the Adam optimizer [25], AlexNet [26] and ResNet [17] architectures, different batchsize, dropout [41], and *L*_2_ weight decay [27], but results were the same and in some cases even worse. Using less pooling layers also yielded worse results. Our model can (over)fit the training data, as evidenced by high training quality metrics and low training loss, Fig. 3F. However, when splitting training data into 5-fold cross-validation, the validation quality metrics and loss change direction after about 40 epochs (Figs. 3A–E). Therefore, we stop after 40 epochs when training on the entire training data in order toavoid overfitting.

**Figure 3:**
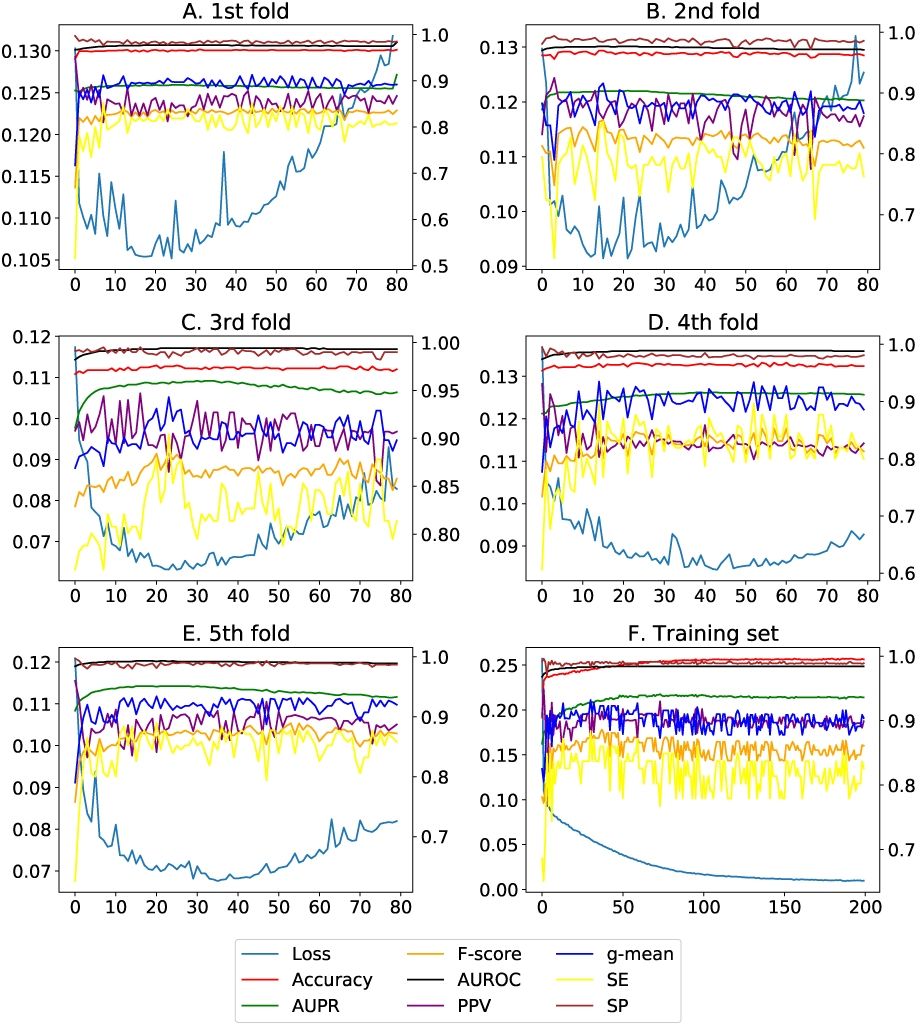
Validation quality metrics for five rounds of crossvalidation on 90% of the human dataset (A–E) and training quality metrics trained on 90% of the human training set (F). Various validation metrics become worse (indicating overfitting) after about 40 epochs, hence we use modelstrained for 40 epochs in evaluations.

## 3. Experiments

### 3.1 Datasets

In our experiments, we used datasets used in [36, 45], which were obtained from miRBase [15], NCBI (https:CODE [7] and snoRNA-LBME-db [30]. The datasets are termed human, cross-species, and new.

#### Algorithm 1 Variable-sized inputs

~~~
1: **function** BACKPROPAGATION(*network, TrainSet*)
2:		*batchSize* ← 1
3:		*count* ← 0
4:		*batchGrad* ← 0
5:		**while** *True* **do**
6:			*input* ← *BatchInput*(*TrainSet, batchSize*)
7:			*label* ← *BatchLabel*(*TrainSet, batchSize*)
8:			*grad* ← *ComputeGradient*(*input, label*)
9:			*batchGrad* ← *batchGrad* + *grad*
10:			*count* ← *count* + 1
11:			**if** *count* = 64 **then**
12:				*UpdateNetworkParameter*(*batchGrad*)
13:				*count* ← 0
14:				*batchGrad* ← 0
~~~

#### Algorithm 2 Fixed-sized inputs

~~~
1: **function** BACKPROPAGATION(*network, TrainSet*)
2:	*batchSize* ← 64
3:	**while** *True* **do**
4:		*input* ← *BatchInput*(*TrainSet, batchSize*)
5:		*label* ← *BatchLabel*(*TrainSet, batchSize*)
6:		*grad* ← *ComputeGradient*(*input, label*)
7:		*UpdateNetworkParameter*(*grad*)
~~~

Table 1 shows the number of positive and negative samples in each of the datasets. In total, there are 3230 positive and 23934 negative samples, which leads to a class imbalance. To cope with the class imbalance, we assigned more weight to positive samples in the cross-entropy loss function.

**Table 1:** Statistics of the three datasets. We have to cope with a class imbalance problem.

|  | human | cross-species | new |
| --- | --- | --- | --- |
| Positive samples | 863 | 1677 | 690 |
| Negative samples | 7422 | 8266 | 8246 |

Fig. 4 shows the histogram of sequence lengths in our dataset. While the mean length of RNA sequences is 96, there are still sequences with more than 300 nucleotides. The longest and shortest RNA sequences consist of 398 and 45 nucleotides, respectively.

**Figure 4:**
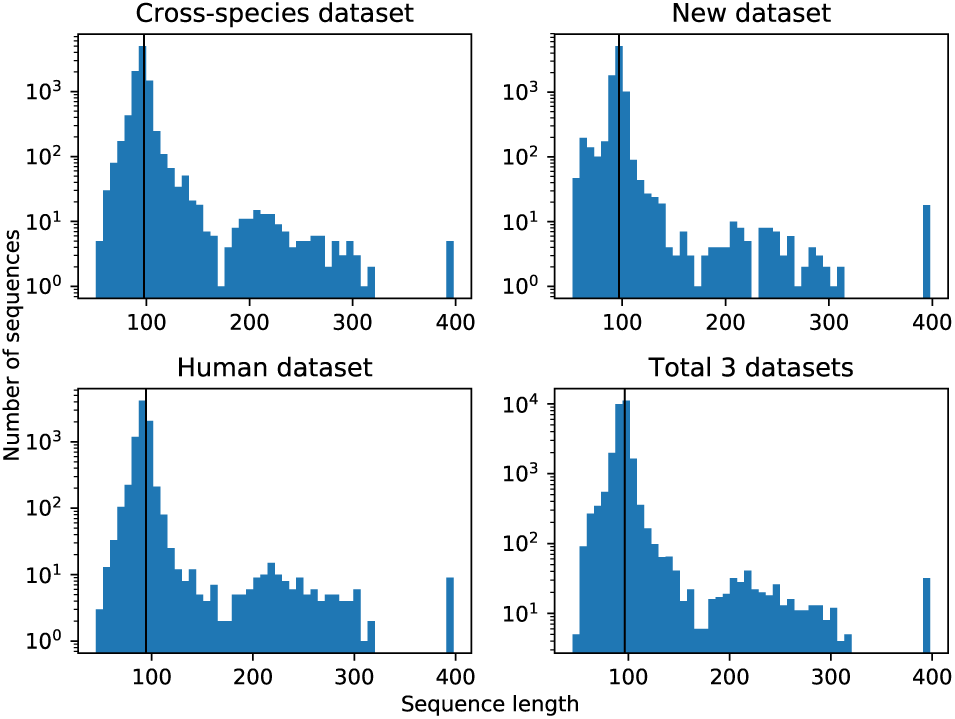
Histogram of length of RNA sequences. Black line is the average length. The sequence length is variable, with the minimal, average and maximal length being 45, 96and 398, respectively.

In our model-building phase, 90% of the human and cross-species datasets are used. We choose model architecture and hyperparameters (i.e. number of epochs, learning rate, batch size) based on 5-fold cross-validation results (cross-validation is run on 90% of the human dataset separately and on 90% of the cross-species dataset separately, but we found aforementioned hyperparameters that worked well for both), which is the standard procedure to tune hyperparameters in pre-miRNA identification literature [45, 36].

After cross-validation, we use three standard evaluation (test) procedures:

1. Train on the aforementioned 90% of the human dataset, test on the remaining 10% of the human dataset, as done in [36].
2. Train on the aforementioned 90% of the cross-species dataset, test on the remaining 10% of the cross-species dataset, as done [36].
3. Train on the entire cross-species dataset, test on the entire new dataset, as Tran et al. [45] proposed.

We have not used the test 10% of the human and cross-species datasets, nor any samples from the new dataset at any point during training nor hyperparameter search.

### 3.2 Cross-validation and test performance

#### 3.2.1 Experiment settings

We test the success of the model against previous work done in the field. As described above, we tested on 10% of human and cross-species datasets, and all of the entire new dataset. We also report results of cross-validation, which is also a common evaluation in literature. For comparison, we used sensitivity (SE), specificity (SP), positive predictive value (PPV), F-score, g-mean, area under the receiver operating characteristic (AUROC), and area under the precision-recall curve (AUPR). Results are shown in Tables 2, 3, 4. Results of miRBoost, CSHMM, triplet-SVM, microPred, MIReNA and deepMiRGene are referred from [36]. We also rerun deepMiRGene with the code the the authors provided, and also included the reproduced results in Tables 2, 3, 4. Formulas for metrics described are caculated as following: TP: Σ true positive, TN: Σtrue negative, FP: Σfalse positive, FN: Σfalse negative, SE = TP / (TP + FN), SP = TN / (TN + FP), PPV (precision) = TP / (TP + FP), F-score = 2TP / (2TP + FP + FN), 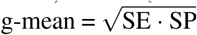. The decision threshold is 0.5.

**Table 2:**
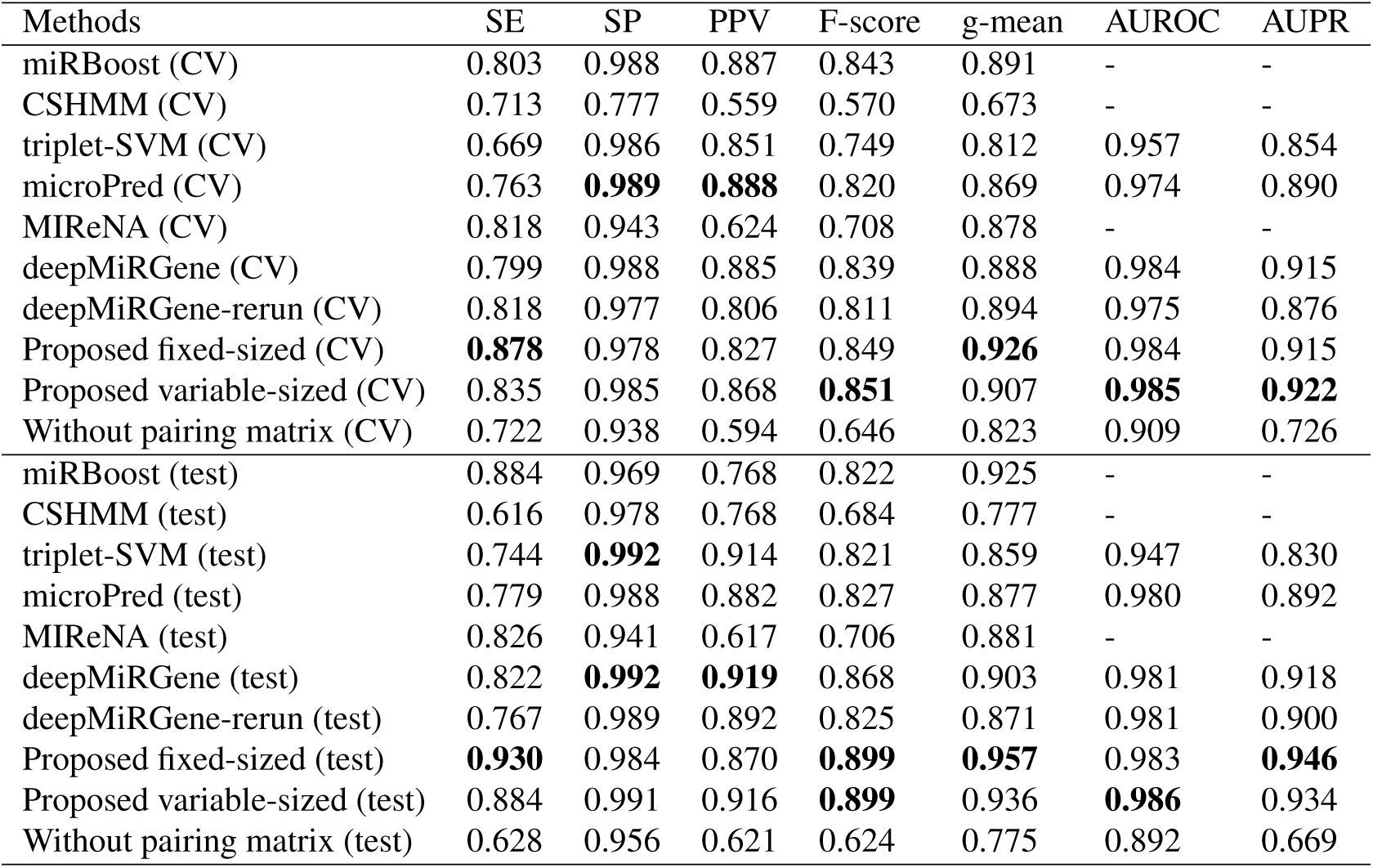
Results on the human dataset. We outperform state-of-the-art methods in 5 out of 7 measurements.

| Methods | SE | SP | PPV | F-score | g-mean | AUROC | AUPR |
| --- | --- | --- | --- | --- | --- | --- | --- |
| miRBoost (CV) | 0.803 | 0.988 | 0.887 | 0.843 | 0.891 | - | - |
| CSHMM (CV) | 0.713 | 0.777 | 0.559 | 0.570 | 0.673 | - | - |
| triplet-SVM (CV) | 0.669 | 0.986 | 0.851 | 0.749 | 0.812 | 0.957 | 0.854 |
| microPred (CV) | 0.763 | **0.989** | **0.888** | 0.820 | 0.869 | 0.974 | 0.890 |
| MIReNA (CV) | 0.818 | 0.943 | 0.624 | 0.708 | 0.878 | - | - |
| deepMiRGene (CV) | 0.799 | 0.988 | 0.885 | 0.839 | 0.888 | 0.984 | 0.915 |
| deepMiRGene-rerun (CV) | 0.818 | 0.977 | 0.806 | 0.811 | 0.894 | 0.975 | 0.876 |
| Proposed fixed-sized (CV) | **0.878** | 0.978 | 0.827 | 0.849 | **0.926** | 0.984 | 0.915 |
| Proposed variable-sized (CV) | 0.835 | 0.985 | 0.868 | **0.851** | 0.907 | **0.985** | **0.922** |
| Without pairing matrix (CV) | 0.722 | 0.938 | 0.594 | 0.646 | 0.823 | 0.909 | 0.726 |
| miRBoost (test) | 0.884 | 0.969 | 0.768 | 0.822 | 0.925 | - | - |
| CSHMM (test) | 0.616 | 0.978 | 0.768 | 0.684 | 0.777 | - | - |
| triplet-SVM (test) | 0.744 | **0.992** | 0.914 | 0.821 | 0.859 | 0.947 | 0.830 |
| microPred (test) | 0.779 | 0.988 | 0.882 | 0.827 | 0.877 | 0.980 | 0.892 |
| MIReNA (test) | 0.826 | 0.941 | 0.617 | 0.706 | 0.881 | - | - |
| deepMiRGene (test) | 0.822 | **0.992** | **0.919** | 0.868 | 0.903 | 0.981 | 0.918 |
| deepMiRGene-rerun (test) | 0.767 | 0.989 | 0.892 | 0.825 | 0.871 | 0.981 | 0.900 |
| Proposed fixed-sized (test) | **0.930** | 0.984 | 0.870 | **0.899** | **0.957** | 0.983 | **0.946** |
| Proposed variable-sized (test) | 0.884 | 0.991 | 0.916 | **0.899** | 0.936 | **0.986** | 0.934 |
| Without pairing matrix (test) | 0.628 | 0.956 | 0.621 | 0.624 | 0.775 | 0.892 | 0.669 |

**Table 3:**
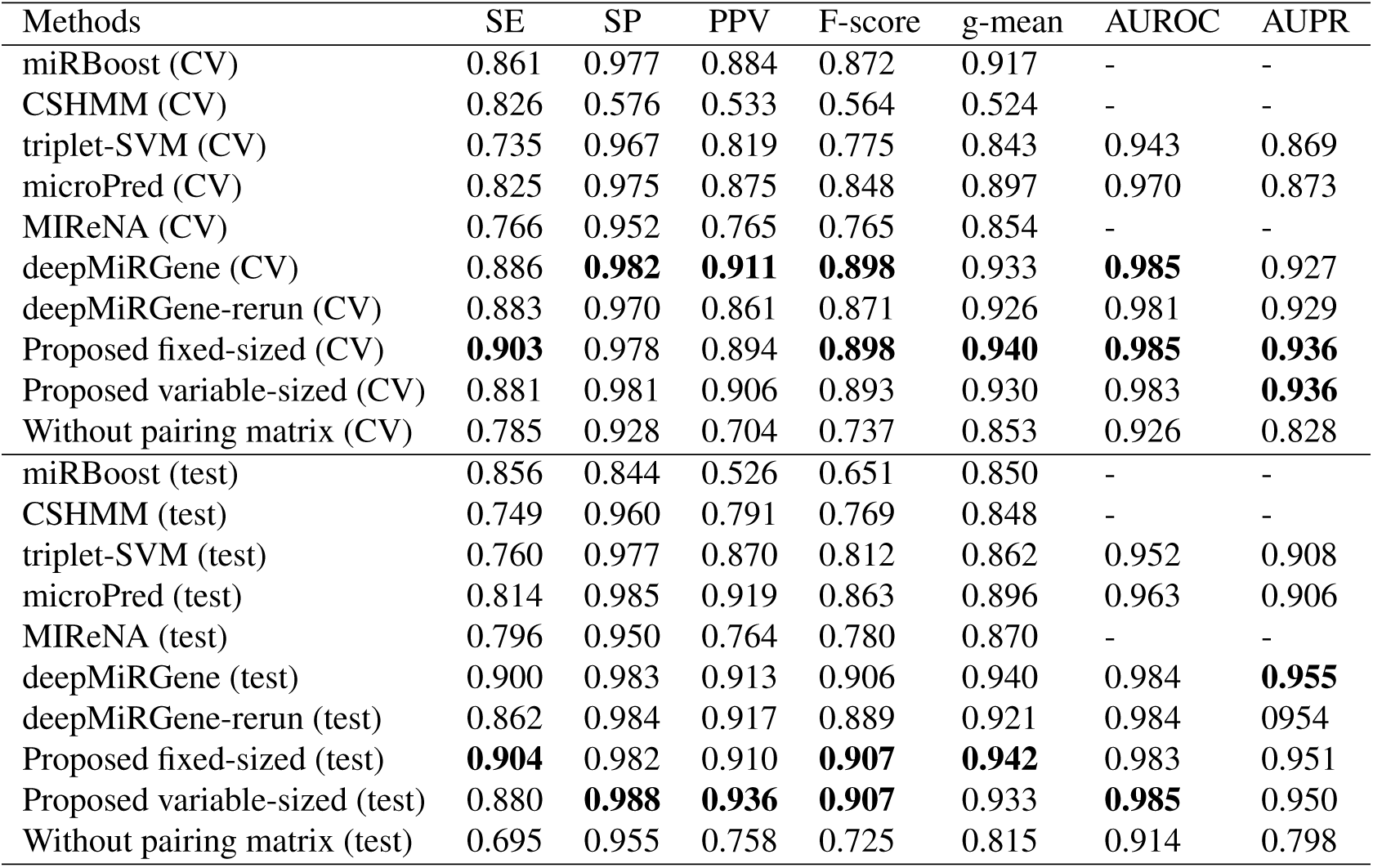
Results on the cross-species dataset. We achieve comparable results with deepMiRGene and outperform other methods.

**Table 4:** Results on the new dataset. We outperform state-of-the-art methods in 4 out of 7 measurements, especially for F-score.

| Methods | SE | SP | PPV | F-score | g-mean | AUROC | AUPR |
| --- | --- | --- | --- | --- | --- | --- | --- |
| miRBoost | **0.921** | 0.936 | 0.609 | 0.733 | 0.928 | - | - |
| CSHMM | 0.536 | 0.069 | 0.046 | 0.085 | 0.192 | - | - |
| triplet-SVM | 0.721 | **0.981** | 0.759 | 0.740 | 0.841 | 0.934 | 0.766 |
| microPred | 0.728 | 0.970 | 0.672 | 0.699 | 0.840 | 0.940 | 0.756 |
| MIReNA | 0.450 | 0.941 | 0.392 | 0.419 | 0.650 | - | - |
| deepMiRGene | 0.917 | 0.964 | 0.682 | 0.782 | 0.941 | **0.981** | 0.808 |
| deepMiRGene-rerun | 0.899 | 0.968 | 0.700 | 0.787 | 0.933 | 0.980 | 0.798 |
| Proposed fixed-sized | 0.917 | 0.967 | 0.696 | 0.792 | **0.942** | 0.979 | **0.864** |
| Proposed variable-sized | 0.859 | **0.981** | **0.779** | **0.817** | 0.918 | 0.979 | 0.818 |
| Without pairing matrix | 0.855 | 0.938 | 0.535 | 0.658 | 0.896 | 0.958 | 0.798 |

Since we are dealing with a skewed dataset (there are alot more negative sequences), AUPR and AUROC are important because they are not susceptible to class imbalance. AUPR is especially important for us as our main goal is to detect the positives. Precision measures the probability for detecting positive samples correctly, and it is not affected by the large number of negative samples in our dataset.

#### 3.2.2 Cross-validation results

Looking at the 5-fold cross-validation results, our approach outperformed state-of-the-art methods on the human dataset and achieved competitive results on the cross-species dataset. Specifically for the human dataset, the sensitivity (0.878) is 6% better than the best of others (0.818), while the F-score, g-mean, AUROC and AUPR are better than others as well. This indicates that our approach can handle imbalanced data, and raises hopes for good results the test phase as well.

#### 3.2.3 Test results

The proposed method outperforms previous methods quite consistently. In the human dataset, the test results of every metric except two (specificity and PPV) are better than all other methods. For the cross-species dataset, our method shows results comparable to other methods in terms of specificity, PPV, and F-score and AUROC. On the new dataset, it achieves the the highest scores in specificity, PPV, F-score and AUPR. Also, in all datasets, we achieve the best F-score (0.899 for human, 0.907 for cross-species, and 0.817 for new). Even in metrics where our methods do not have the best score, it is very close to the best in almost all the cases. In the human and new datasets, we can see that our method gives us an improvement over deepMiR-Gene.

We can also see how deep learning approaches without handcrafted features, namely deepMiRGene and our model, perform compared to previous machine learning methods. The new methods have better results in the human dataset in every metric except for specificity, where triplet-SVM gives the same score as deepMiRGene. For both CV and test parts of cross-species, deep learning approaches perform better than all of the previous methods. For the new dataset, miRBoost gives the best result in terms of sensitivity, and triplet-SVM has the same specificity as our proposed model, but in every other category, deep learning approaches have better results. Considering all of the datasets and results, deep learning methods outperform the machine learning solutions.

When comparing deepMiRGene with our method, our approach gives better results in all test datasets. For the human dataset, our approach gives improvements in every metric except for specificity and PPV. For the cross-species dataset, our approach gives better results in terms of every metric other than AUPR. Lastly, for the new dataset, our model performs better in every metric except for AUROC. The results when training on the cross-species dataset and testing on new dataset (Table 4) demonstrate that our approach identifies pre-miRNAs in a new species well although it is trained on a mixed species dataset.

It is also important to compare our two approaches with each other. Without the pairing matrix, the results are much worse in each dataset and each metric. Therefore, the pairing matrix is an important part of our method. Fixed-sized and variable-sized inputs give similar results for most of the categories. With fixed-sized inputs, we need more memory to train than with variable-sized inputs. Therefore, we encourage using variable-sized inputs for this task.

Considering all the results and comparisons with other methods, the results show that our proposed method gives the best performance in identifying pre-miRNAs.

## 4. Discussion and conclusions

The results when using the pairing matrix are much better than just using the input sequence. It means the pairing matrix and secondary structure play an important role in miRNA identification. By including secondary structure information as input, the feature space becomes bigger. Therefore, we need a big dataset for good data generalization. In addition to small numbers of samples, the classes are also imbalanced. A promising direction for future work to improve identification would be to collect more data and then to use bigger neural networks such as AlexNet [26] or ResNet [17].

We proposed to use a “disentangled” representation of the RNA secondary structure, namely the pairing matrix, as input to a 2D convolutional neural network architecture to extract features automatically. We obtain state-of-the-art results, especially for F-score, on the three benchmark datasets human, cross-species and new. In the future, we will try to expand our method to other tasks related to miRNA such as miRNA target prediction and miRNA function prediction [39].

